# The salmon louse genome: copepod features and parasitic adaptations

**DOI:** 10.1101/2021.03.15.435234

**Authors:** Rasmus Skern-Mauritzen, Ketil Malde, Christiane Eichner, Michael Dondrup, Tomasz Furmanek, Francois Besnier, Anna Zofia Komisarczuk, Michael Nuhn, Sussie Dalvin, Rolf B. Edvardsen, Sven Klages, Bruno Huettel, Kurt Stueber, Sindre Grotmol, Egil Karlsbakk, Paul Kersey, Jong S. Leong, Kevin A. Glover, Richard Reinhardt, Sigbjørn Lien, Inge Jonassen, Ben F. Koop, Frank Nilsen

## Abstract

Copepods encompass a range of ecological roles from parasites to phytoplankton grazers linking primary producers to higher trophic levels. Despite these important roles, copepod genome assemblies are scarce. *Lepeophtheirus salmonis* is an economically and ecologically important ectoparasitic copepod. We present the 695.4 Mbp *L. salmonis* genome assembly containing ≈60% repetitive regions and 13081 annotated protein-coding genes. The genome comprises 14 autosomes and a ZZ-ZW sex chromosome system. Assembly assessment identified 92.4% of the expected arthropod genes. Transcriptomics validated annotation and revealed a marked shift in gene expression after host attachment, including downregulation of genes related to circadian rhythm coinciding with abandoning diurnal migration. The genome shows evolutionary signatures including loss of peroxisomes, numerous FNII domains, and an incomplete heme homeostasis pathway suggesting heme proteins to be obtained from the host. Despite large capacity to develop resistance against chemical treatments *L. salmonis* exhibits low numbers of many genes involved in detoxification.

## 1 Introduction

Genomes may be seen as sensitive cellular organs that monitor and respond to environmental challenges [1], and consequently increasing our understanding of genomes will improve our understanding of the harboring organisms, and *vice versa*. Yet, extracting meaning from genetic code has proven challenging, and the large increase in the availability of genomes has raised as many questions as answers [2, 3]. One of the reasons for this is that while genomes constrain the range of possible phenotypes, they do not necessarily determine the phenotypes within that range [4, 5]. This implies that interpretation of phenotype-genotype relationships requires biological data and sequence information from numerous species representing a wide range of environmental backgrounds and evolutionary lineages. The most valuable data will therefore come from taxa or ecological adaptations with previous scarce coverage. Copepods are aquatic arthropods with central ecological roles as grazers linking primary producers to higher trophic levels [6], vectors for disease [7] and parasitic pathogens [8]. Despite their widespread importance, only few copepod genome assemblies have been published, and annotated genomes are, with the exception of *Caligus rogercresseyi* [9], restricted to free-living species [9-12].

Copepods date back to the Cambrian Period, and have since then diversified into numerous lineages, including several types of fish parasites such as the ectoparasitic sea lice (family Caligidae). Sea lice are parasites that spend most of their life on their hosts and are expected to have evolved physiological and behavioral adaptations to their host habitat, including the host’s immune defenses [13, 14]. Although host-parasite adaptations go both ways, sea lice can be deleterious to the host, as exemplified by the effects of *Lepeophtheirus salmonis, Caligus elongatus,* and *C. rogercresseyi* on salmonids [15–19]. While the *Caligus* species typically are generalists infecting many unrelated host species, *L. salmonis* depends exclusively on salmonid hosts for successful reproduction. Such host-parasite relationships are commonly manifested through functional adaptations reflected in the genomes [20, 21]. Hence a genome assembly from a host specialist such as *L. salmonis* would be valuable for comparison to the recently published genome of the host generalist *C. rogercresseyi* [9].

A reliable genome assembly for *L. salmonis* will also be of instrumental value in its own right. Since its inception in the 1970’s, intensive salmon farming has dramatically increased host density, which facilitates salmon lice transmission and population growth [22]. This is problematic for both wild and farmed salmonids since salmon lice cause lesions when feeding, which can entail reduced growth, osmotic imbalance, secondary infections and increased mortality [8, 23–25]. *L. salmonis* has a direct transmission cycle without intermediate hosts allowing it to respond directly to the abundance of hosts [8]. The life cycle consists of eight stages separated by molts [26, 27]. Adult males fertilize adult females which carry the eggs until they hatch into planktonic nauplius larvae. The planktonic larvae pass through two molts before reaching the infective copepodid stage. The remaining 5 stages (chalimus I and II, preadult I and II and adults) are parasitic on the host. Adult females may live for more than 452 days [28] and continuously produce egg strings containing hundreds of eggs [29]. This life history strategy assures high fecundity and wide copepodid dispersal and may be regarded as an adaptation to historical low densities of the anadromous hosts. Salmon farming has driven *L. salmonis* population increases that in turn has resulted in need for salmon louse population control. This has until recently been achieved using chemotherapeutics, but resistances have appeared and spread repeatedly [30–32] leading to a shift towards mechanical delousing strategies, such as low pressure washing and warm baths, which significantly compromise animal welfare [33]. Salmon louse infestations therefore remain a main obstacle for sustainable salmon farming, representing a threat to wild salmonid stocks and causing annual losses estimated to be around one billion EUR [29]. There has therefore been an instrumental need for a high-quality salmon louse genome assembly for studies of basal biology, studies uncovering resistance mechanisms, and studies aiming at vaccine development.

Here, we present the annotated LSalAtl2s genome assembly of the Atlantic subspecies of salmon louse, *Lepeophtheirus salmonis salmonis* [34] which has proven to be a valuable tool for exploring genome evolution, gene regulation and gene function in this highly adapted parasite. The genome may prove particularly valuable in conjunction with the genome of its hosts, e.g. Atlantic salmon [35]. Our analysis expands the current knowledge of genome diversity in arthropods in general and in copepods in particular. It also reveals a set of features reflecting the parasitic lifestyle, including loss of protein families, loss or reduction of metabolic pathways, and even loss of the entire peroxisome organelle.

## 2 Materials and Methods

### 2.1 Sequencing, assembly and annotation

The Ls1a strain [28] of *L. salmonis salmonis* inbred for 27 generations was sequenced to 181-fold assembly coverage in a hybrid approach using Illumina, 454 life sciences and Sanger sequencing to facilitate construction of a *de novo* assembly (see Supplementary Material section S1 for details). Several experimental assemblies of the genomic sequence data were constructed, and after an evaluation process (see Supplementary Material section S2), the final assembly process was decided on. To produce the final scaffolded LSalAtl2s assembly, the 454 life sciences reads were mapped to the salmon louse mitochondrial genome [36] using BWA [37], and matching reads were removed. The remaining reads were assembled using Newbler [38], version 2.6, using the `-large` option. In order to adjust for homopolymer errors that are common artifacts of the 454 pyrosequencing process [38], the Illumina reads were mapped to the contigs and a new consensus sequence was produced using samtools mpileup [39] to collect mapping information, bcftools view `-cg` - to generate per position variant information, and vcfutils.pl vcf2fq [40] to call the consensus assembly. The assembly was scaffolded by SSPACE [41] in a series of iterations using libraries with increasing read pair distances (paired reads with 260 bps distance, then paired reads with 500 bps distance and finally mate pair reads with 3-6 kbps distance). All scaffolding was performed with parameters `-k 3 -a 0.7`, except for Illumina mate pair data, where the parameters were `-k 5 -a 0.3`. The scaffolds were aligned to the SwissProt and UniProt90 databases [42], and scaffolds that were found to contain genes with bacterial annotation, and with no RNA-seq mapping *and* fragmented/incomplete DNA mapping were regarded as contamination and removed. Finally, scaffolds had terminal N’s removed and were filtered for length, removing all scaffolds with fewer than 200 nucleotides with mapped Illumina genomic DNA (gDNA) reads and all scaffolds shorter than 500 nucleotides without mapped Illumina gDNA reads.

Protein-coding gene models were constructed using Maker (v2.27). Firstly, a *de novo* repeat library was generated using the program RepeatModeler (v1.0.5), which was subsequently used by Maker to mask repetitive regions of the genome. To enhance gene predictions, transcriptome data derived from the inbred salmon louse strain was used. Samples for Illumina sequencing were derived from all stages; unfertilized eggs, early developing eggs (pooled eggs obtained 0-24 hours after fertilization), late developing eggs (pooled eggs obtained 2-7 days after fertilization), nauplius I, nauplius II, copepodids, chalimus I and II, preadult I and II females, preadult I and II males, adult females and adult males. Resulting RNA-seq data were mapped to the genome with Tophat (v1.3.2) and assembled using Cufflinks (v1.1.0) to provide supporting evidence for the gene build. Expressed Sequence Tag (EST) data from the parasitic copepods *L. salmonis salmonis*, *Caligus clemensi*, *C. rogercresseyi* and *Lernaeocera branchialis* [43] were used by Maker in its est2genome prediction mode to predict an initial set of genes, which was then used, in conjunction with all known *Daphnia pulex* proteins, to train the gene finder SNAP. SNAP was run on the salmon louse genome and retrained with the resulting genes. The salmon louse EST set was used to train the gene finder Augustus (v2.5.5). Maker was then run on the repeat masked sequence using the trained SNAP and Augustus programs, the EST alignment data, and all protein sequences from the phylum Arthropoda available from the UniProt Knowledgebase (downloaded 17. May 2013). A second gene set was derived by running Maker on the genome without prior repeat masking. InterProScan 5 (RC7) was run to identify protein domains and to map GeneOntology terms to salmon louse genes. The final gene set comprises all genes from the first run, together with genes from the second run containing InterPro domains and not overlapping with genes from the first run. The Benchmarking Universal Single-Copy Orthologs (BUSCO) V 5.0.0 datasets (downloaded 22. February 2021) for Arthropoda, Metazoa, and Eukarya (arthropoda_odb10, metazoa_odb10, eukarya_odb10) were used to check for presence of genes that are expected to be conserved across the included lineages. OrthoMCL (v2.0.8) was used to compare the gene set with other species. Homology relationships between salmon louse genes and genes from other species were detected using the Ensembl Compara Gene Trees pipeline [44]. KEGG Orthology assignments and KEGG pathway maps were generated by submitting the Ensembl predicted protein sequences to the KEGG Automatic Annotation Server (KAAS) [45].

### 2.2 Repeat analysis

Comparative repeat analysis of crustaceans was done as follows: repeat families were modelled *de novo* from each genome assembly using RepeatModeler 2.0.1 [46] with the pipeline extensions for classification of LTR elements enabled. Then, RepeatMasker version 4.1.1 [47] was run in sensitive mode using the generated repeat families, both programs were used with rmblastn version 2.10.0+. For salmon lice, repeat families were generated based on the *L. salmonis salmonis* LSalAtl2s assembly only, these were used for all four salmon louse assemblies. The following assemblies were used: *Acartia tonsa* - GCA_900241095.1 [11], *Caligus rogercresseyi* - GCA_013387185.1 [9], *Daphnia pulex* - GCA_900092285.2 [48], *Eurytemora affinis* - GCF_000591075.1 (I5K initiative, unpublished), *L. salmonis salmonis* – LSalAtl2s (this manuscript), *L. salmonis* female - GCA_001005205.1 (Leong et al., unpublished), *L. salmonis* male - GCA_001005235.1 (Leong et al., unpublished), *Leopeophtheirus salmonis onchorhynchi* (Pacific subspecies of *L. salmonis* [34]) - GCA_000181255.2 (GiLS, unpublished), *Oithona nana* - GCA_900157175.1 [10], *Tigriopus californicus* - GCA_007210705.1 (Scripps Institution of Oceanography, unpublished), *Tigriopus kingsejongensis* - GCA_012959195.1 [12] and *T. kingsejongensis* [49].

A recent unpublished copepod assembly found in GenBank (*L. salmonis onchorhynchi* - GCA_016086655.1) was excluded from whole genome analyses in compliance with the responsibilities for data users set forth in the Fort Lauderdale Agreement, Section C.2 [50].

### 2.3 Genomic structure and recombination

Using data from a related project sequencing salmon lice from multiple locations in the Northern Atlantic Ocean [30], Single Nucleotide Polymorphism (SNP) markers were identified and subsequently used to construct a linkage map. The data consisted of 5098 SNP markers genotyped on 12 full sib families [30, 51], each consisting of two parents and 46 offspring. Both genotype and pedigree data were handled in LepMAP3 [52]. The data was first checked and corrected for erroneous or missing genotypes with the ParentCall2 function. SNPs were then assigned to chromosomes with the SeparateChromosomes2 function. The default parameter of SeparateChromosomes2 lead to 13 autosomes, but previous work based on a larger number of markers [53], showed that the genome of *L. salmonis* comprises 14 autosomes. The SeparateChromosomes2 function was therefor run again with a higher threshold parameter (`LODlimit=15`), identifying 14 autosomes and one sex chromosome. Finally, SNP order and sex specific recombination distances were estimated on each chromosome separately by using the OrderMarkers2 function with default parameters.

### 2.4 Expression analysis of an Spo11 endonuclease ortholog

Triplicate pairs of testes and ovaries from adult inbred Ls1a males and females were dissected and placed into RNAlater, extracted using Qiagen RNeasy kits and cDNA was made using Affinity script (Agilent) and random hexamer primers. Quantitative real time PCR (qPCR) on dilution series was run using target (mRNA) *LsSpo11* and control (mRNA) *LseEF1α* assays in triplicate 10 µl reactions (for details see Supplementary Material section S6). The PCR efficiencies of the *LsSpo11*and *LseEF1α* [54] qPCR assays were comparable in the entire dilution range. The assays were further evaluated for sample or assay specific trends in efficiencies (eg. induced by PCR inhibitors) observing no such trends. Comparative analysis of six gonad samples (three testes and three ovaries) was conducted in triplicate reactions on a three step dilution series. Relative gene expression was calculated using the established *LseEF1α* standard gene [54] and ovary as calibrator tissue according to the ∆ΔC_T_ method [55]. C_T_ values diverging by more than 0.5 cycles from the average of triplicate reactions were omitted and the 95% confidence intervals were calculated using average ΔC_T_ values for each of the triplicate reactions.

### 2.5 Phylogenomic analysis

Based on the Ensembl Compara analysis, 173 1:1 orthologs were identified of the species included in the analysis. These protein sequences were used to create a concatenated sequence from each species that were aligned using default settings in ClustalX [56]. This alignment was used for phylogenetic inference using MrBayes [57] implementing a GTR + I + Γ model, run with 4 simultaneous chains for 1 million generations. Full names of the included species: *Ciona intestinals*, *Danio rerio*, *Homo sapiens*, *Mus musculus*, *Schistosoma mansoni*, *Trichinella spiralis*, *Brugia malayi*, *Caenorhabditis elegans*, *Ixodes scapularis*, *D. pulex*, *L. salmonis*, *Pediculus humanus*, *Tribolium castaneum*, *Drosophila melanogaster*, *Aedes aegypti* and *Anopheles gambiae*.

### 2.6 Gene presence and absence

Annotations for all included species (Fig. 3) were retrieved from Ensembl (release 101) and Ensembl Metazoa (release 48) using BioMart either manually or via the R package biomaRt [58]. Protein families and domain expansions and deletions were assessed based on the *L. salmonis* InterProScan annotations compared to sixteen existing genome annotations stored in Ensembl (see Supplementary_Table_Compara_Domains). Pathway analysis was based on KEGG pathways [59]. Gene presence and absence calls were based on reciprocal best BlastP hits between the predicted *L. salmonis* proteome and UniProtKB or GenBank, requiring a best hit within the same group of orthologs or EC-number, but not necessarily the same species. To call a sequence likely absent, we also required absence of representative InterProScan hits and performed reciprocal TBlastN/BlastX searches between the LSalAtl2s assembly and UniProtKB/GenBank. Blast searches were performed using the NCBI Blast suite versions 2.6.0+ and 2.9.0+ [60, 61] with E-value threshold < 1E-6. Additional analytical steps and accessions of queries used in Blast searches are described in the Supplementary Material.

### 2.7 Transcriptomic analysis of antennae and gut for annotation validation

For transcriptome analysis of different tissues, lice from Ls1a strain, also used for gDNA sequencing, were utilized. Ovaries, testes, and intestine were dissected from adult lice. Antennae were sampled from adult females, adult males and copepodids. Copepodid antennae were taken from planktonic copepodites. Legs were dissected from both male and female adult lice. For a more detailed description see Supplementary Material section S3. In addition to the stage samples described above, attached copepodids were also sampled each day from one to six days after infection. For RNA sequencing, purified RNA from copepodids was combined from two following sampling days each. RNA was extracted and purified as described in Supplementary Material section S4. Due to low RNA quantity in the adult female antenna sample, this RNA was amplified, using the SeqPlex RNA Amplification Kit (Sigma Aldrich). The following libraries were generated: HBR-1: ovaries, pooled sample; HBR-2: testis, pooled sample; HBR-3: legs adult male and female pooled; HBR-4: antenna adult male, pooled sample; HBR-5: antenna copepodid, pooled sample; HBR-6: Antenna adult female, pooled sample; HBR-7: copepodids sampled 1 and 2 days post infection at 10 °C; HBR-8: copepodites sampled 3 and 4 days post infection at 10 °C; HBR-9: copepodids sampled 5 and 6 days post infection at 10 °C; HBR-10: intestine, adult female, pooled sample.

Library preparation and sequencing was conducted by Fasteris SA (Geneva, Switzerland). A more detailed description is available in Supplementary Material section S3. Libraries were prepared for 50 bp single-end reads and were sequenced in multiplexed mode on Illumina HiSeq 2000 with up 10 million reads per sample. Illumina’s stranded RNA-seq protocol (TruSeq, with polyA selection) with forward direction primer were used for all libraries, except for library HBR-4N which was prepared using total RNA-protocol with normalization by Duplex-specific Nuclease (DSN) due to low RNA concentration. Data from library HBR-6 is not strand specific due to the amplification process applied.

RNA-sequences were aligned and counted with respect to the Ensembl genome annotation. Samples were quality clipped and eventual adapter sequences removed using TRIMMOMATIC [62]. Trimmed sequences were aligned against the LSalAtl2s assembly with the splice-aware aligner STAR [63] using the Ensembl transcript models for constructing the genome build. Read counts were obtained for all transcripts in the Ensembl annotation using the Bioconductor package easyRNAseq [64]. Library-size normalized read counts were calculated using the Bioconductor package edgeR [65] and are given in counts per million (CPM). Further details on methods used for creating Fig. 1 are given in Supplementary Material section S3.

**Figure 1.**
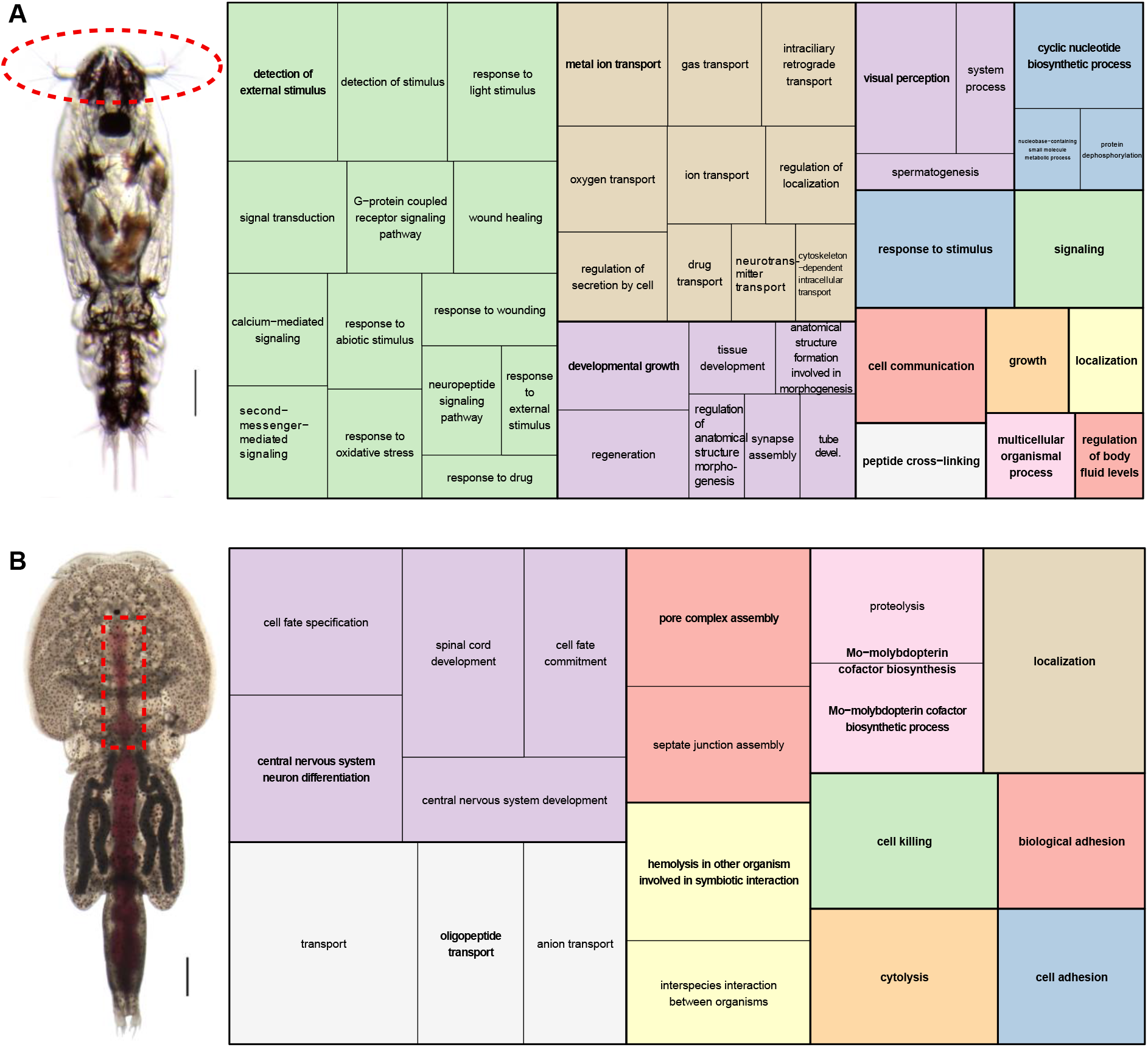
GO Enrichment analysis of transcripts expressed in A) copepodid antenna, B) adult intestine (anterior). The approximate dissection areas are indicated by red dashed lines on the photographs of the respective stages to the left: A) planktonic copepodid, scale bar 0.1mm; B) adult female with blood-filled intestine, scale bar 1mm. Significantly enriched GO-terms from the BP ontology are depicted in the form of a tree-map to the right, with clusters of semantically similar terms indicated by map colors. Centroids are typeset in bold. Note, GO-terms related to the central nervous system in B) may be caused by inclusion of ganglia that is localized close to the anterior intestinal lumen.

### 2.8 Transcriptomic analysis of transition from planktonic to attached lifestyle

For planktonic *L. salmonis* copepodids, published data were used [66]. Briefly, these copepodids of the LsGulen strain were hatched from three different egg-string pairs on second day after molting to the copepodid stage. To obtain attached copepodids, farmed Atlantic salmon reared at 10 °C were infected with copepodids from LsGulen strain [28]. Experimental procedures were performed in accordance with Norwegian animal welfare legislation (permit ID7704, no 2010/245410). Lice were sampled from six different fish (sextuple samples) at one, three, and five days after infection. For details see Supplementary Material section S12. Library preparation, RNA sequencing, and data processing were executed as previously described for planktonic copepodids [66]. Sequences were aligned to the LSalAtl2s assembly using STAR and transcript counts were obtained using featureCounts [67]. Differential gene expression analysis was implemented by DESEq2 using Galaxy [68]. An adjusted p-value of 0.05 was taken as cut off.

## 3 Results and discussion

### 3.1 Assembly and annotation

The final LSalAtl2s assembly has a size of 695.4 Mbp distributed among 36095 scaffolds with a scaffold N50 of 478 Kbp (contig N50: 6044). Annotation results in prediction of 13081 protein coding and 482 non-coding RNA genes. The BUSCO datasets for Arthropoda, Metazoa, and Eukaryota were used to assess completeness of the genome by checking for presence of genes that are expected to be conserved across these taxa. Among the genes in the BUSCO V 5.0.0 Arthorpoda dataset 92.4% were complete (90.4% as single copy genes, 2.0% as duplicated genes) while 4.4 % of the BUSCO genes were missing and 3.2% were fragmented (results for Metazoa and Eukaryota are shown in Supplementary_Table_GenomeStats). Previous transcriptomics studies that used LSalAtl2s as reference genome also reported high average mapping rates for RNA-sequencing reads (>93%) and high rates of unique mapping to a single genomic location (>88%) when using a highly sensitive splicing-aware aligner [66, 69].

We used transcriptomic data to validate the gene annotations by functionally characterizing copepodid antenna and adult intestine *L. salmonis* tissues by Gene Ontology (GO) terms: (Fig. 1). For copepodid antenna, “detection and response to stimulus” as well as “signaling” were the main GO-terms. For the adult intestine a more variable distribution of GO-terms was found, including “Proteolysis and hemolysis in other organism involved in symbiotic interaction” and ”transport”. Through this approach, we confirmed the sensory function of antennae and the digestive function of the intestine, thus lending support to the soundness of the annotation.

Genome statistics for the salmon louse and selected organisms illustrate that the salmon louse has a relatively compact genome with an unexceptional number of predicted genes (Table 1). Copepod genome sizes may reach extreme values of 32 pg (≈ 32 Gb) haploid DNA per cell as found in *Paraeuchaeta norvegica* [70]. Of the examined species, the genome of the free-living copepod *T. californicus* is more compact than the salmon louse with comparable gene content while both *T. kingsejongensis* and *C. rogercresseyi* are reported to have smaller genomes and around twice as many genes (Table 1).

**Table 1.** Comparison of genome metrics of selected ecdysozoan species and representative assemblies. The LSalAtl2s assembly has N50 comparable to assemblies based on Sanger sequencing and generally higher N50 than pure Illumina-based assemblies. Assembly names, depth and method information was taken from GenBank or the associated publication if not available in GenBank. Scaffold N50 was computed if not found in GenBank, GC%, and Repeat% were computed from the genome sequence. Repeat content is given in percent of total bases masked by RepeatMasker. Species are (from left): Lepeophtheirus salmonis, Caligus rogercresseyi, Tigriopus kingsejongensis, Tigriopus californicus, Acartia tonsa, Pediculus humanus, Caenorhabditis elegans. An extended table with additional species, information and accession numbers is available (Supplementary_Table_GenomeStats).

| Species | *L. sal.* | *C. rog.#* | *T. kin.* | *T. cal.* | *A. ton.* | *P. hum.* | *C. ele.* |
| --- | --- | --- | --- | --- | --- | --- | --- |
| Assembly Mb | 695.4 | 478.2 | 338.6 | 191.1 | 989 | 110.8 | 100.3 |
| Assembly level | Scaffold | Chromosome | Scaffold | Chromosome | Scaffold | Scaffold | Chromosome |
| Sequencing technology | Illumina HiSeq 2000, 454, Sanger | PacBio, Illumina Hiseq4000 | PacBio | Illumina, PacBio | Illumina | Sanger | Sanger |
| Depth | 175 | 155 | 70 | 180 | 50 | 8.5 | NA |
| #Scaffolds | 36 095 | 21 | 938 | 459 | 351 850 | 1 882 | 7 |
| Contig N50 | 6 044 | 38 017 | NA | 44 438 | 3 244 | 34 097 | NA |
| Scaffold N50 Kbp | 478 | 27 803 | 1 473 | 15 806 | 4 | 497 | 17 494 |
| GC% | 31 | 34 | 47 | 42 | 32 | 27 | 35 |
| Repeat % | 60* | 57* | 25* | 28* | 45* | 20 | 18* |
| # genes | 13 085 | 25 399 | 25 470 | 15 577 | NA | 10 785 | 20 191 |
| BUSCO Arthropoda | C:92.4%, F:3.2% | C:61.0%, F:10.9% | C:95.5%, F:2.3% | C:93.1%, F:3.6% | C:57.8%, F:24.6% | C:97.1%, F:2.3% | NA |

\*De novo RepeatModeler run was used to first generate repeat families and then RepeatMasker was run with the output. In *P. humanus*, RepeatMasker was used with the public DFAM repeat library and taxon information.
#Numbers are taken from GenBank; slight deviations were found between these and the values in [9].

### 3.2 Extensive repetitive elements in L. salmonis

*De-novo* detection by RepeatModeler yielded 4076 unique repeat families in the LSalAtl2s assembly. For comparison, we included genome assemblies from seven other aquatic crustaceans, as well as all four public assemblies of *L. salmonis* (Supplementary Material Fig. S4-1). Despite differences in sequencing depth, assembly software, and fragmentation, both GC-content and the proportion of repeat families is very similar between all four *L. salmonis sp.* assemblies providing confidence that the observed repeat frequency is not affected by assembly parameters. For all assemblies analyzed, the majority of repeats could not be assigned to any known family (Supplementary Material section S4 and Fig. S4-1).

The total proportion of masked sequences was ≈60% in the *L. salmonis salmonis* LSalAtl2s assembly (61% - 62,5% in assemblies of *L. salmonis onchorhynchi*, see Supplementary Material section S4). This is the highest repeat content of all sequenced crustacean genomes presently available. The second highest repeat content is found in the only other available Caligid genome presently available; *C. rogercresseyi* with 57% repeats. It should be noted that Gallardo-Escárate [9] reported a lower *C. rogercresseyi* repeat content (51.9%), possibly due to differences in parameter settings (e.g. running RepeatMasker in sensitive mode). Caligid repeat content is among the highest in all sequenced arthropod genomes. Notwithstanding recent advances in methods for repeat detection, Caligid repeat content is also comparable to much larger genomes, for example the Atlantic salmon (*Salmo salar,* 58–60%, 2.97 Gb [35]), locust (*Locusta migratoria*, ≈60%, 6.5 Gb, [71]), and Axolotl, the largest sequenced animal genome (*Ambystoma mexicanu,* 65.6%, 18-30Gb [72]).

Evidently, *C. rogercresseyi* and *L. salmonis spp.* are rich in autonomous mobile genetic elements, DNA transposons and retro-elements, and these represent much larger fractions of the genome than in the sequenced free-living crustaceans. The classifiable Transposable Elements (TEs) with highest copy number in *L. salmonis* belong to the Tc1 and Mariner families of transposons (Supplementary Material section S4 and Supplementary_data_TableS4-1.gz). These families are extant in various copy-numbers in eukaryote lineages, and their expansion in *L. salmonis* might have a beneficial effect on genome plasticity and evolution [73, 74]. There is growing evidence for horizontal transposon transfer (HTT) between different species, including fish and their parasites [75–77] and future studies are needed to establish whether some caligid TEs are active or have been interchanged with their host via HTT.

### 3.3 Genomic structure and recombination

A linkage map was constructed based on 5098 previously identified SNP markers [30]. Of these markers, 5062 were assigned to linkage groups. Using Blat, 4786 markers were mapped unambiguously to 1250 scaffolds with a total size of 534 Mbp (77% of the assembly). 4127 markers mapped to scaffolds with markers from only one linkage group. Of these, 398 scaffolds contained only one marker, and 777 scaffolds contained >2 markers from the same linkage group (in total 3729 markers). 75 scaffolds with a total size of 72 Mbp (10.4% of the genome assembly) contained markers from multiple linkage groups (659 markers). These results suggest that synteny of approximately 10% of the assembly may be affected by errors in assembly, scaffolding or linkage group assignment.

Linkage map analysis showed that the *L. salmonis salmonis* genome comprises 15 linkage groups (LGs) ranging in size from 3 to 157 cM (Fig. 2). A total of 462 Mbp (66% of the 695.5 Mbp assembly) was assigned to linkage groups through scaffolds without ambiguous LG assignment. LG12 exhibited extremely low recombination rates in both sexes while the remaining LGs exhibit frequent recombination in males and extremely low recombination rates in females wherefore marker distances were calculated from a male map (Fig. 2). A lack of heterozygous markers was observed in LG15 in females, whereas heterozygosity was common in males for the same LG (Fig. 2B) confirming that LG15 is a sex chromosome as previously suggested [53]. These observations functionally explain the 1:1 male:female ratio in salmon lice and are compatible with both a ZZ-ZW and a ZZ-Z0 sex chromosome configuration and alternative chromosome configurations with several sex chromosomes as previously reported in both deuterostome and protostome animals [78, 79]. As only female lice were sequenced the W chromosome is not included in the assembly.

**Figure 2.**
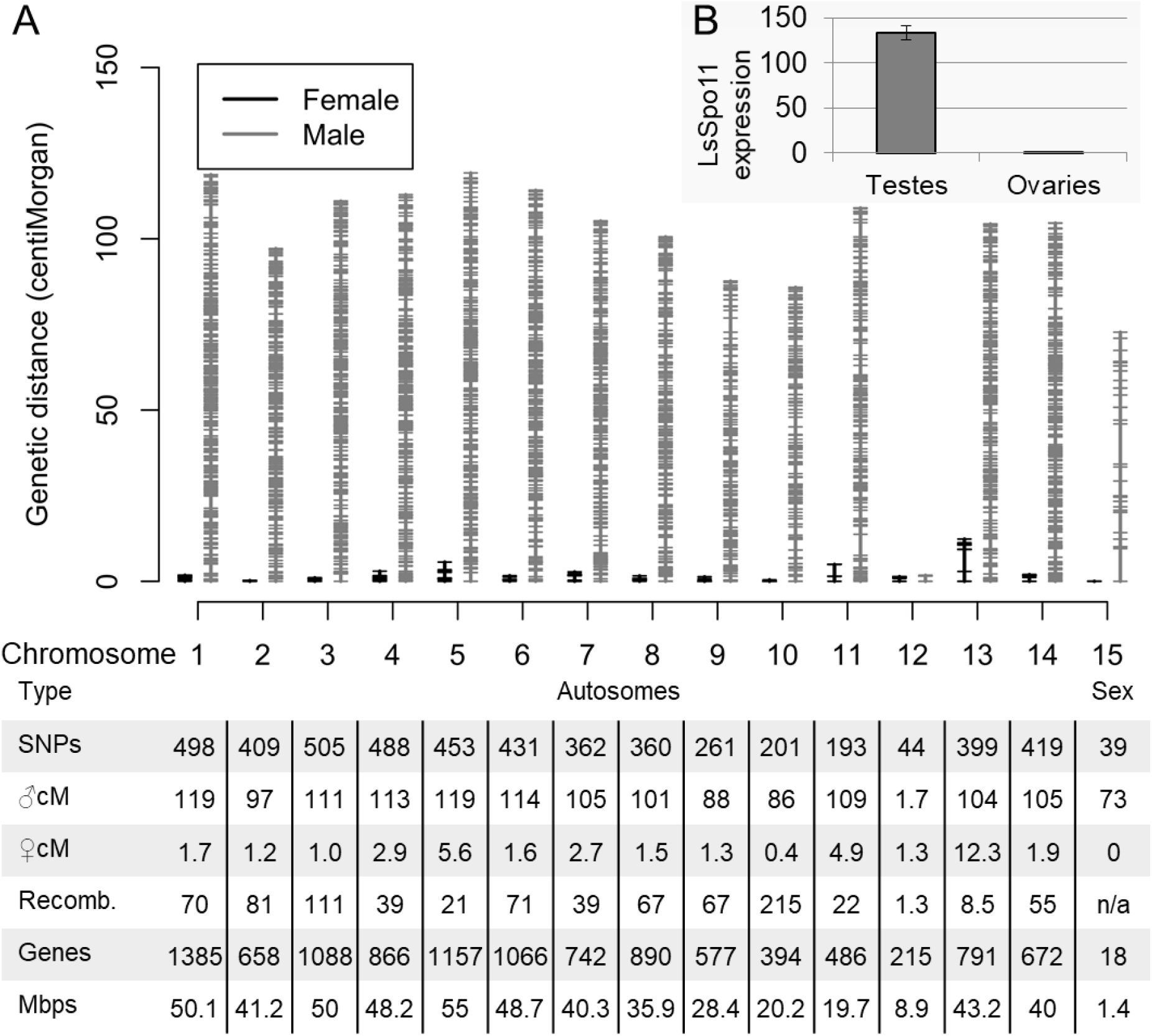
Linkage map, chromosome statistics and LsSpo11 expression. A: Linkage map showing the relative size of male and female linkage groups and marker distribution. The table shows chromosome type for each LG, the number of SNP markers included in the map, male and female chromosome sizes in centimorgan (cM), the male/female recombination ratio (Recomb.), the number of genes located to the individual LGs, and the number of base pairs comprised in unambiguously assigned contigs (total base pairs comprised in assigned scaffolds indicated in brackets). Note that the total number of genes and base pairs assigned to LGs are lower than the total figures for the assembly as not all scaffolds are assigned to a LG. B: Relative expression of LsSpo11, involved in double stranded break formation during recombination.

Independent sequencing of an Atlantic *L. salmonis* male (NCBI sequence read archive SRX976782) allowed independent mapping of male and female reads to the assembly. This showed that the majority of scaffolds received average mapping of both male and female reads indicating them to be autosomal (Supplementary Material Fig. S5-1). A smaller proportion of scaffolds received average mapping of male reads and 0.5x average mapping of female reads indicating them as a Z-type sex chromosome (present in two copies in males and a single copy in females). All of the scaffolds assigned to LG15, save one, fell in this category indicating LG15 to be a Z chromosome. Finally, a number of scaffolds received no mapping of male reads and 0.5x average mapping of female reads indicating them as belonging to a W-type sex chromosome (present as single copy in females only). This group contained no scaffolds assigned to LGs as markers not present in both sexes were omitted from the LG-analysis. In summary *L. salmonis salmonis* is heterogametic with a ZW type chromosome system possibly comprising more Z and/or W chromosomes. A single scaffold assigned to LG15 received average mapping from both males and females suggesting it to represent a homologous region shared between the Z and W chromosomes. These findings agree with the conclusions drawn by [53].

The salmon louse is heterochiasmic as genetic recombination is predominantly found in the homogametic males. As the topoisomerase-like Spo11 protein is involved in genetic recombination by inducing double stranded breaks [80, 81] and is present as a single copy gene in the salmon louse genome, we investigated the possible involvement of the LsSpo11 protein in the recombination profile observed. The *LsSpo11* gene is highly expressed in testes compared to ovaries with a difference exceeding 100-fold (Fig. 2B). These results are in keeping with the anticipated function of LsSpo11 in recombination and may explain the almost absent recombination in females. Although recombination in only one sex is unusual, a similar pattern is found in other species including *D. melanogaster* that has XX-XY type sex chromosomes, two regular chromosomes that recombine in the females only and a small “dot” chromosome that does not recombine in either sex [82–85]. Interestingly, in similarity to the *D. melanogaster* ‘dot’ chromosome, the salmon louse also has a chromosome sheltered from recombination in both sexes; the chromosome corresponding to LG12 (Fig. 2 and [53]).

### 3.4 Phylogenomic analysis

The official gene set predicted by Ensembl comprised 13081 coding genes, and among these the Ensembl Compara pipeline identified 173 1:1 orthologs across 16 selected species that were used in a whole genome phylogenetic reconstruction (Fig. 3). *L. salmonis* groups together with *D. pulex* and these two “crustacean” species occur as a sister group to the included insects where the tick (*I. scapularis*) represents the most basal arthropod branch of the included species (Fig. 3). The observed grouping pattern is in line with previous studies [86, 87]. The included parasites have fewer genes (14285 genes in average) compared to the non-parasites (19708 genes in average) which could suggest a general gene loss in the parasite species. Similar observations have been found in bacteria [88] and it could be hypothesized that parasites may require fewer genes than their free-living relatives since parasites can exploit their host for pre-processed nutrients, metabolic intermediates, etc. However, parasites are also likely to have additional requirements related to parasite-host interactions and it should be considered that phylogenetic bias may influence the interpretation. Hence comprehensive studies with additional taxa should be conducted before firm conclusions regarding trends in gene numbers and parasitism are made.

**Figure 3.**
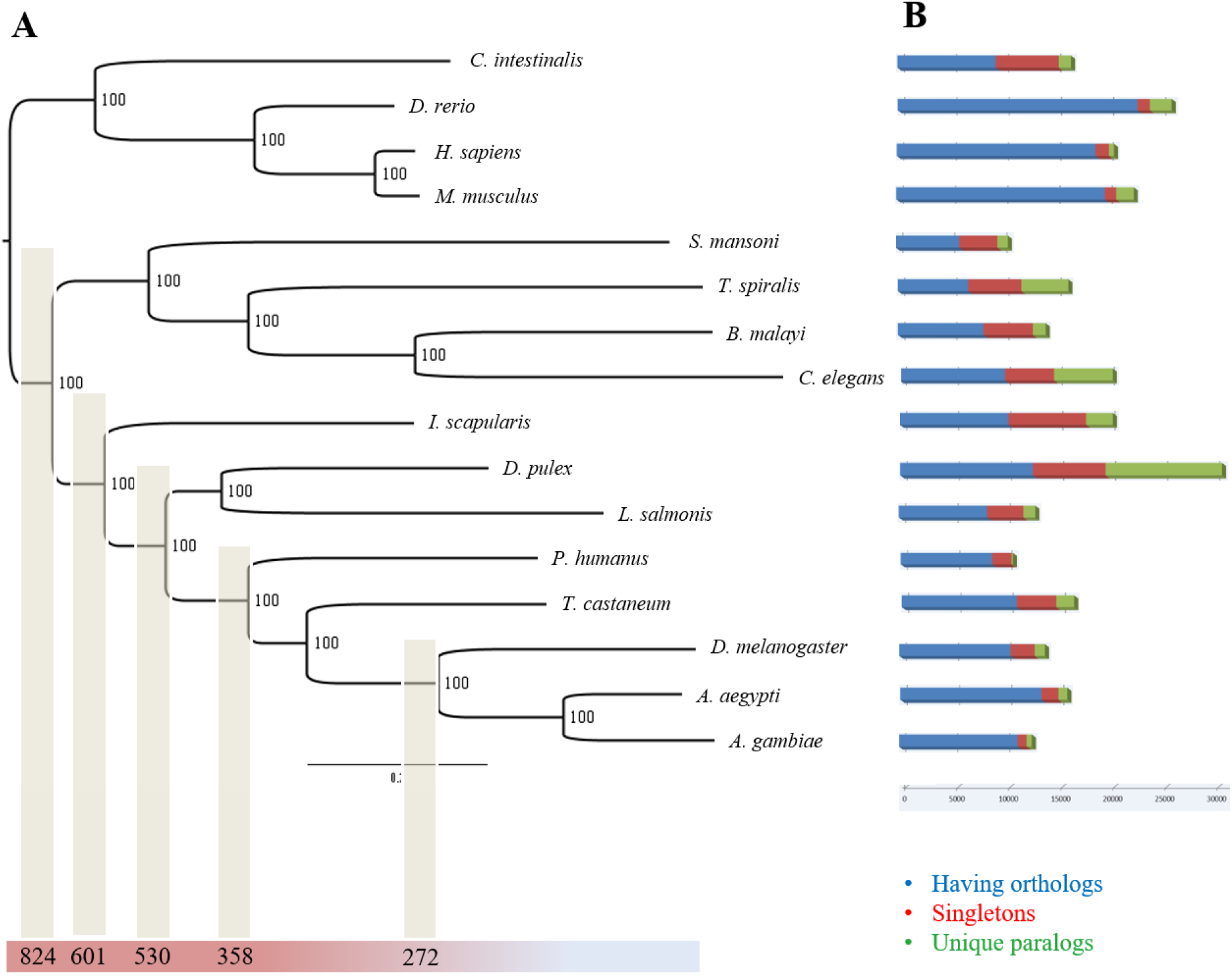
Whole genome phylogeny based on 173 1:1 orthologous genes (A) and the number of genes in each species having orthologs, singletons and unique paralogs (B). The number on the branches represents posterior probabilities and all showed 100% support for the given branching pattern. The divergence time (million years) is given below some of the main lineages and is based on estimates from TimeTree [89]. Full names of the included species: Ciona intestinals, Danio rerio, Homo sapiens, Mus musculus, Schistosoma mansoni, Trichinella spiralis, Brugia malayi, Caenorhabditis elegans, Ixodes scapularis, Daphnia pulex, Lepeophtheirus salmonis, Pediculus humanus, Tribolium castaneum, Drosophila melanogaster, Aedes aegypti and Anopheles gambiae.

### 3.5 Gene losses and expansions

The gene presence and absence analysis revealed some protein groups and pathway elements to be absent, and others to be found in surprisingly high or low numbers. A comprehensive overview can be found in Supplementary_Table_Compara_Genes. Among the most conspicuous findings are: A complete lack of annexins, a large expansion of SHK domains, an incomplete heme homeostasis pathway, loss of the genes needed to sustain peroxisomes, a reduction in most genes expected to be involved in detoxification, and a large number of FNII domains - a domain previously considered to be vertebrate specific. These findings are treated in further detail below.

#### 3.5.1 L. salmonis lacks annexins

Annexins are present in most organisms and are involved in a range of basic biological processes such as calcium metabolism, cell adhesion, growth and differentiation. Cantacessi *et al.* [90] assessed 35 species of invertebrates for annexins finding these in all but the nematode *T. spiralis* and the mollusc *Oncomelania hupensis*. Surprisingly, annexin domains (PF00191) were not found in the *L. salmonis* genome and could not be detected in the *C. rogercresseyi* (GCA_013387185) or *E. affinis* (GCF_000591075) genomes either. Annexin was, however, found in the *T. kingsejongensis* genome showing that its absence is not a shared copepod trait.

#### 3.5.2 L. salmonis has a large expansion of the SHK domains (PF01549)

The SHK domain was first identified in a potassium channel inhibitor from a sea anemone [91, 92]. A total of 125 SHK domains in 24 genes were identified in *L. salmonis* which is an extreme number for any arthropod species. *T. kingsejongensis* has only two SHK domains and the cladoceran *D. pulex* has 10 SHK-domains whereas some nematodes have a large number of SHK-domains (Supplementary_Table_Compara_Domains). It has been shown that SHK binds to potassium channels in human B and T lymphocytes and prevents activation of these [93]. We speculate that the SHK-domain containing proteins in the salmon louse could be important for immunomodulation of the host and warranting further investigation.

#### 3.5.3 Incomplete heme homeostasis pathway

Host blood is an important part of the salmon louse diet and is rich in nutrients, among them proteins, lipids, and trace metals. Heme is an iron-containing macrocycle and an essential prosthetic group in aerobic life and its biosynthesis involves a highly conserved 8-step enzymatic pathway. It has recently been shown that 7 of the 8 genes coding for this pathway are absent from the genome of the salmon louse. Instead, it contains an intestinally expressed gene coding for a putative heme receptor linked to heme absorption [94]. Furthermore, the genome encodes genes for several requisite heme-binding proteins such as cytochrome c and a recently characterized heme-peroxidase [95] and the mitochondrial and nuclear genomes contain all components of the mitochondrial electron transfer chain. Considering that excess heme is accessible from salmon blood, we conclude that the salmon louse is a natural heme auxotroph (see Supplementary Material section S7 for details). This feature combined with the initial free-living stages of the lifecycle, calls for further investigation into the mechanisms of heme trafficking and storage, in particular within oocytes. Homologous loci related to heme-metabolism, -binding and -trafficking detected in the genome are listed in the Supplementary Material file Supplementary_Table_Heme. Taking into account heme and porphyrin toxicity (reviewed in [96]), the absence of any known detoxification mechanism is surprising. The classical conserved two step pathway via heme oxigenase and biliverdin-reductase is lacking from the genome unlike in other arthropods. Other mechanisms of heme detoxification such as formation of hemozoin or retaining heme in the peritrophic matrix, as described for hematophagous insects [97], currently lacks evidence in the salmon louse. Interestingly, out of the eight conserved steps of heme biosynthesis the salmon louse genome contains only the enzyme responsible for one: Coproporphyrinogen oxidase (*Cpox*, EMLSAG08964). In humans, mutations in genes encoding heme biosynthesis, including *Cpox*, can cause autosomal dominant porphyria via accumulation of toxic intermediates of porphyrins (reviewed by [98]). The preservation of CPOX in the salmon louse may indicate a role in clearing such compounds. Finally, since *L. salmonis* lacks Heme oxygenase (*Hmox*) known to be responsible for Heme degradation, we hypothesize that the *L. salmonis* has a hitherto unknown heme detoxification pathway or - more generally - resistance mechanism against heme toxicity.

#### 3.5.4 L. salmonis lack peroxisomes

Peroxisomes are organelles with a common evolutionary origin and conserved traits in biogenesis [99]. Despite their shared phylogenetic origin, they are involved in metabolic processes that may diverge significantly between species and tissues [100, 101]. Four Pex genes are conserved over a wide taxonomic range and may be considered marker genes required for presence of peroxisomes: *Pex3*, *Pex10*, *Pex12* and *Pex19* [102]. A fifth *Pex* gene with bacterial orthologs, *Pex5*, is also ubiquitously present in peroxisome containing organisms [102]. InterProScan analysis indicated that none of the 5 core *Pex* genes were present in *L. salmonis*. To reassert that the core *Pex* genes were indeed missing in the salmon louse, the LSalAtls2s assembly was also scrutinized by Blast analysis using a large number of *Pex* genes as query (Supplementary Material section S6). Absence of the Pex core genes show that the salmon louse is unable to sustain peroxisomes [102]. To investigate if lack of peroxisomes is a conserved feature among copepods, we searched available genome assemblies for *C. rogercresseyi, Calanus finmarchicus, T. californicus and E. affinis* (Supplementary Material section S6). We found that both *T. californicus* (order Harpactgocoida), *E. affinis* and *C. finmarchicus* (order Calanoida) possess all core elements of peroxisomes while the fish louse *C. rogercresseyi* (order Siphonostomatoida, same family as *L. salmonis*) lack core *Pex* genes (Supplementary Material section S6). This indicates that the lack of peroxisomes may be an adaptation to parasitism among members of the family Caligida, such as *L. salmonis* and *C. rogercresseyi*, or maybe even shared among members of order Siphonostomatoida in general as suggested by the lack of core PEX protein encoding ESTs from other siphonostomatoids presently represented in GenBank (*L. branchialis*; 14927 ESTs, and *C. clemensi;* 46858 ESTs). While peroxisomes are generally considered ubiquitous organelles, their absence has previously been reported in various taxa, including parasitic platyhelminths and nematodes, and even the free-living appendicularian *Oikopleura dioica* [103]. A key function of peroxisomes is to facilitate catalase mediated reduction of reactive H_2_O_2_ to oxygen and water. This capacity does not appear to be lost as a single catalase gene is retained and expressed in the salmon louse (see licebase.org; EMLSAG00000007315). This gene is, unsurprisingly, devoid of PTS1/2 peroxisome targeting signal motifs (cf. Islinger et al. [101]).

#### 3.5.5 Peptidases in L. salmonis

We compared different types of peptidase domains across the included species (Supplementary Material section S9) with particular reference to species that are blood feeding. For most species including *L. salmonis*, serine peptidases (e.g. Trypsin PF00089) form the most abundant peptidase domain. However, there are some striking differences between the different species where the tick deviates from the other included blood feeders by having a much larger proportion of M13 peptidases (139 N-terminal domains and 106 C-terminal domains). Ticks have intracellular digestion and the high proportion of M13 peptidases could be a signature for this property [104]. On the other hand, the salmon louse has the highest proportion of M12A (astacin) peptidase domains (67 domains in total) of the five species included in the InterProScan analysis (see Supplementary_Table_Compara_Domains and Supplementary Material Figure S9-1). This is more than twice as many as what is found in *D. pulex*.

#### 3.5.6 Detoxification and stress-response

The salmon louse possesses a remarkable capacity for adaptation to most chemical delousing agents [30–32], including hydrogen peroxide [105, 106]. We therefore hypothesized ac expansion of gene-families related to xenobiotics. Interestingly, the opposite is the case. When comparing the frequency of gene-families with putative xenobiotics related roles to other arthropods and vertebrate species in Ensembl, three important gene-families are strongly reduced while only one is slightly expanded with a total of four members (Supplementary_Table_Detox).

Using the LSalAtl2s assembly and additional transcriptome data, Humble *et al.*, [107] reported that the family of cytochrome P450 (*Cyp*) genes is the most compact compared of all arthropods. Besides their role in detoxification of various substrates, enzymes of the Cyp family contribute, among others, to metabolism of steroids, fatty acids, and vitamins. Notably, all *L. salmonis Cyp* genes fall into sub-family class E, group I (IPR002401), and none is classified as class E, group II (containing major insect *Cyp* genes with detoxification ability) by the InterProScan analysis.

The second group of significantly underrepresented families are transporters of the ATP-Binding Cassette type (ABC). The ABC transporter family in *L. salmonis* has been extensively surveyed [108].

The third compacted gene family in the salmon louse are the Glutathione S-transferases (GSTs); one of the most important families of detoxifying enzymes in nature (reviewed by Oakley [109]). There is growing evidence that GSTs are induced by and may contribute to the clearing of synthetic pyrethroids such as cypermethrin in terrestrial arthropods [110] and aquatic crustaceans [111]. Amelioration of pyrethroid toxicity by GSTs may be due to their antioxidant capacity [112]. Despite their importance, the salmon louse genome carries only 7 and 13 hits to the conserved C- and N-terminal GST domains, respectively, compared to 55 hits to the N-terminal domain (IPR004045) in *C. elegans*. Again, this is the lowest number of GSTs in the arthropod genomes in our comparison.

We found only one expanded gene-family that might be related to detoxification. We identified four paralogs of the major vault protein (MVP), which is so far un-reported in crustaceans and hexapods. We further detected single copy orthologs in other copepods (*C. finmarchicus, T. kingsejongensis* and *T. californicus*, see Supplementary_Table_Vault_Genes and Supplementary Material section S10). MVP is highly conserved throughout many eukaryote linages and the major constituent of vaults, the largest cytoplasmic ribonucleoprotein complex in eukaryotes known. We detected orthologs of the minor vault proteins Tep1, and several Parp-like orthologs. (Supplementary_Table_Vault_Genes). The less conserved ribonucleotide components, vault-associated RNA (vtRNA), could not be detected.

The function of vault particles is still elusive, but MVP and other vault components have been implicated in a variety of functions, such as signaling pathways, regulation of apoptosis, autophagy, inflammation, nuclear-cytoplasmic transport, and multi-drug resistance (MDR) in cancer [113, 114]. In a recent study on *L. salmonis*, *Mvp* transcription was upregulated together with *Cyp* genes and other stress responsive genes under cypermethrin exposure [115]. Mammalian MVP is induced by or responds to biotic and abiotic stressors and toxins in vivo and vitro [116–118]. Unlike most eukaryote genomes which harbor either none or a single copy of the *Mvp* gene, *L. salmonis* contains four paralogous Mvp-like sequences, likely stemming from gene duplication events. Implication in resistance and stress-response render MVP and vaults interesting for future studies in sea lice. However, it should be noted that contribution to MDR is debated (reviewed by Park [119]) and that inferences from cancer cells and mammalian models to an aquatic parasite should be made with caution. Therefore, more studies are needed to elucidate a potential connection of vault duplication to lifestyle or detoxification.

#### 3.5.7 Large expansion of FN2-domains

Fibronectin II domains (FNII-domain, PF00040), commonly considered to be vertebrate specific, are – surprisingly - the largest of the expanded *L. salmonis* families with 192 domain copies. FNII domains are present in 74 annotated genes, mostly alone but regularly in combination with other domains, mainly trypsin (see [120] and Supplementary Material section S11). While FNII domains have not previously been reported in other arthropods, they are described in tubularians [121] and a few other invertebrate groups (see Supplementary Material Table S11-1). Invertebrates commonly have Kringle domains (PF00051) that have been suggested to be ancestral to FNII domains [122]. In addition to FNII domains, we also identified five different proteins containing a single Kringle domain in combination with other domains in *L. salmonis. T. californicus*, belonging to a different order than *L. salmonis*, also has proteins with both Kringle and FNII domains indicating that the divergence of Kringle and FNII domains occurred earlier than previously suggested and probably in a common metazoan ancestor.

While the exact functions of genes with FNII-domains in *L. salmonis* are unknown, the expression profile for all transcripts containing FNII domains in the RNA sequencing samples (Supplementary Material Fig. S11-2) reveals that these genes are expressed at different stages and different tissues, demonstrating that the FNII-domain is widely used in the salmon louse proteome.

#### 3.5.8 Reduced diversity in chemosensory molecules in the salmon louse

It has previously been shown that salmon lice respond to chemical cues from their host [123] and chemical sensing has been proposed to be important for host identification [124]. In Arthropods, chemical sensing is mediated by gustatory receptors (GRs), odorant receptors (ORs), and ionotropic receptors (IRs) and we assessed the genome for orthologs of these proteins. No GRs or ORs were detected in the salmon louse genome whereas 26 IRs were identified (see Supplementary Material section S12) and some play a crucial role in host recognition by copepodids [125]. In the Crustacea *D. pulex* genome 58 GRs were detected [126]. Furthermore, we identified nine putative GRs in *T. californicus* and two in *T. kingsejongensis* whereas no GRs could confidently be identified in *C. rogercresseyi* (see Supplementary Material section S12). This indicates that the loss of GRs may be a signature of Caligidae or Siphonostomatoidea. As expected, many genes are present both *T. kingsejongensis* and *C. rogercresseyi* that contained the PF00060 and PF10613 domains (i.e. iGluR, see Supplementary Material section S12). The more conserved co-receptors (i.e. IR25a, IR8 and IR93a) were detected in both species and for *C. rogercresseyi* five specific IRs (i.e. IR328, IR329, IR333, IR337, IR339) were also found that have orthologs in *L. salmonis*. We could not detect two IR8 in *Tigriopus* or *Caligus*, indicating that a IR8 duplication could be a *Lepeophtheirus* specific feature. However, due to the divergent nature of the IRs all sequences should be experimentally validated before firm conclusions are made (see Table S12-1 in Supplementary_Table_IR).

### 3.6 Gene expression during transition to a parasitic lifestyle

The infective copepodid stage can be investigated in either its non-parasitic planktonic or host bound parasitic phase. By assessing the transcriptome of planktonic versus attached copepodids we aimed to compare these different lifestyles. Gene expression of planktonic copepodids was compared to the expression of parasitic copepodids collected one, three or five days after attachment. Copepodids from the different groups differ in their overall gene expression (Supplementary Material section S13). We found 6273 differentially expressed genes, 3205 transcripts that were up-regulated between planktonic and attached lice consistently in all three comparisons and 3068 that were consistently down-regulated (Supplementary Material section S13). The most relevant GO-terms affected by this transition are shown in Fig. 4. These can be summarized as follows: after attaching to the host, metabolism increases and sensory abilities decrease (all enriched GO-terms can be found in the Supplementary Material section S13). Planktonic copepodids do not feed but rely on internal stores for survival until they find a host [8, 29]. The attachment, and by that the access to food, induces a switch in nourishment and triggers the start to growth and development. Sensing-related genes are strongly downegulated after attachment, while cell cycle, protein synthesis and other core metabolic functions are enriched (Fig. 4). Lipid metabolism is also down regulated after attachment presumably due to the change of nutrition from stored lipids [127] to ingested host tissues. Furthermore, several copies of a gene with Blast hit to the protein Timeless [128], known for regulating circadian rhythm in *Drosophila*, are among the down regulated transcripts. Among the downregulated genes we also find three known for regulating circadian rhythm in *Drosophila*; Shab, Shaw and Cycle [129, 130]. This may reflect the transformation from being a planktonic organism with diurnal migration [131] to a parasitic lifestyle. There are also several downregulated muscle related genes (Myosin, Troponin, Tropomyosin, Actin, Titin and the regulator of muscle contraction Twitchin [132]), suggesting a shift to a more sessile live style. The upregulated genes in attached copepodids include ribosomal proteins, proteosome proteins, transcription and translation factors. The ionotropic receptors probably involved in host recognition [125] are down regulated, while genes with FNII domains (see above) are among the most up regulated genes in parasitic copepodids.

**Figure 4.**
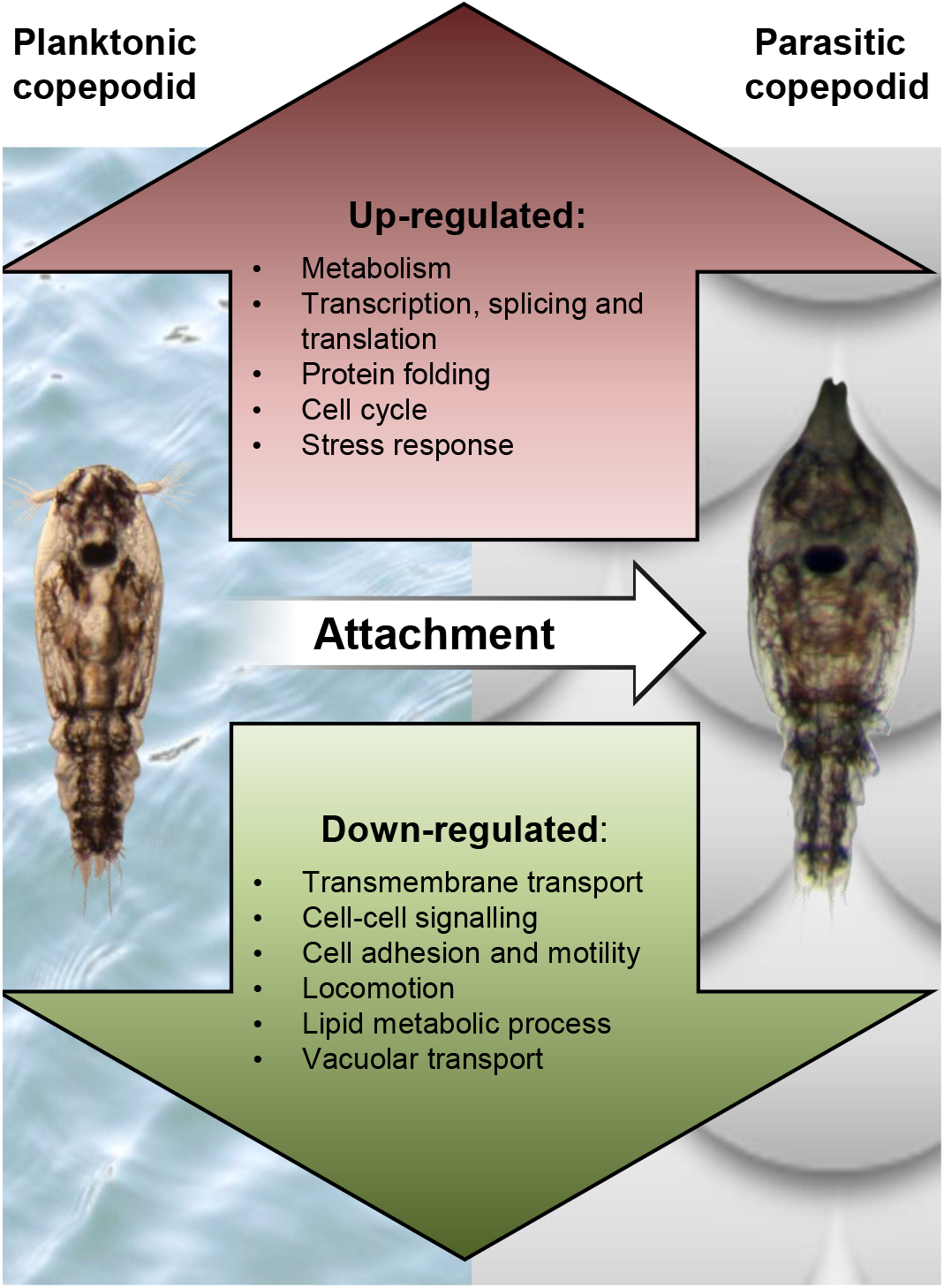
Main biological processes regulated in copepodids after attachment. Gene ontology enrichment in the up- and downregulated genes in copepodids after infecting a host.

## 4 Conclusions

The LSalAtl2s draft genome assembly in conjunction with the gene models in Ensembl Metazoa is the first fully annotated genome of a parasitic copepod and represents an invaluable tool for studies of copepods in general, and the salmon louse in particular. The genome assembly sequence and annotation have been validated by linkage analysis and RNA-sequencing and are reasonably complete with respect to the coding gene inventory. Many genes have already been curated or experimentally validated. The genome is useful as a reference for future genomics and transcriptomics studies, and together with the genome of one of its mains hosts, the Atlantic salmon, it may facilitate investigations into host-parasite interactions. Taken together, the Atlantic salmon louse genome provides a useful tool for further investigations into the genome biology and evolution of copepods, an important crustacean group that is still only sparsely covered by genome assemblies.

## 5 Data Availability

Sequencing raw-data and transcriptomics data have been deposited in SRA and are available in GenBank under BioProjects with accession nos. PRJNA705827, PRJNA413461 and PRJNA577842. The LSalAtl2s scaffold-level assembly and genome annotation including gene models and InterProScan and Compara results are available at Ensembl Metazoa (https://metazoa.ensembl.org/Lepeophtheirus_salmonis). An updated annotation and associated RNA-interference data are kept in LiceBase (https://licebase.org). Original output files from the comparative repeats analyses of various organisms and the BUSCO analysis are available upon request.

## 6 Acknowledgements

We are thankful to Dr. E. Birney (European Bioinformatics Institute) and O. Torrissen (Institute of Marine Research) for their assistance and enthusiasm. The project was funded by The Ministry of Trade and Fisheries, The Norwegian Seafood Research Fund (project 900400) and the Research Council Norway, SFI-Sea Lice Research Centre, grant number 203513/ O30.

